# A singular PpaA/AerR-like protein in *Rhodospirillum rubrum* rules beyond the boundaries of photosynthesis in response to the intracellular redox state

**DOI:** 10.1101/2023.07.13.548891

**Authors:** Manuel S. Godoy, Santiago R. de Miguel, M. Auxiliadora Prieto

## Abstract

Photosynthesis (PS) in Purple Non-Sulphur Bacteria (PNSB) is strictly controlled to avoid energy consumption and to prevent oxidative stress when oxygen is present. PpsR, a transcriptional repressor codified in the Photosynthetic Gene Cluster (PGC), is one of the main players in the regulation of PS related genes. In some cases, PpsR do not sense the environmental cue by itself but delegates this task to another protein like PpaA/AerR anti-repressors, which can ultimately affect PpsR affinity to some promoter regions. In this work, the effects of locus Rru_A0625 product (HP1) on PS regulation of *Rhodospirillum rubrum*, were studied by mutation/complementation and transcriptomic strategies. Rru_A0625 is located next to *ppsR* gene, just like other PpaA/AerR members, and its deletion annuls pigment synthesis in dark micro/anaerobic growth conditions. HP1 shows similarity to PpaA/AerR anti-repressors family, although it does not possess their typical cobalamin binding domain. A transcriptomic analysis of Rru_A0625 deletion mutant showed that HP1 not only has effects on bacterioclorophyll and carotenoid biosynthesis, but also many other biological processes in the cell. The most notorious is the impact on the transcription of the nitrogenase complex components and some accessory proteins. Our results suggest that this new member of the PpaA/AerR family has evolved losing the canonical cobalamin binding domain, but not the redox sensing capability, conserving a not fully understood mechanism of PS regulation.

**IMPORTANCE:** *Rhodospirillum rubrum* vast metabolic versatility places it as a remarkable model bacterium and an excellent biotechnological chassis. The key component of PS studied in this work (HP1) stands out among the other members of PpaA/AerR anti-repressors family since it lacks the most conserved motif they all share: the cobalamin B-12 binding motif. Even with a minimized and barely recognizable aminoacidic sequence, HP1 stills controls PS as the other members of the family, allowing a fast response to changes in the redox state of the cell. This work also shows that HP1 absence affects genes form relevant biological processes other than PS, including nitrogen fixation and stress response. From a biotechnological perspective, HP1 could be manipulated in approaches where PS is not necessary, such hydrogen or polyhydroxyalkanoates production, to suppress unnecessary energy burden.

## INTRODUCTION

One of the key requirements for survival in nature is having a tight and fine-tuned control of energy expenditure. Food and nutrients are generally scarce resources that must be administrated in a balanced way considering the conditions of the moment and anticipating possible future scarcity. Living organisms count on complex sensing systems that allow them to make accurate diagnoses on the availability of resources, in order to process this information and ensure survival by reacting appropriately [1].

In PNSB, these complex sensing-response systems are permanently fraught given their versatile metabolism and the resulting multiplicity of responses they can execute. Their ability to do photosynthesis (PS) [1], fix CO_2_ [2]–[4] and N_2_ [5], ferment sugars and volatile acids [6], extract energy from inorganic sources, such as H_2_ [7] or carbon monoxide [8], demonstrate their metabolic ductility. It also shows the necessity of counting on a complex regulatory system to integrate and coordinate these physiological processes both harmonically and efficiently.

In different PNSB, PS is controlled mainly by three regulatory systems: the two-component system RegBA (also called PrrBA) [9] the anaerobic activator FnrL [10]–[12] and the aerobic repressor PpsR <u>(p</u>hoto<u>p</u>igment <u>s</u>uppression [13], which, contrary to the first two, is exclusively involved in PS regulation. Under aerobic conditions, PpsR binds to a consensus target sequence (TGTN_12_ACA) in the promoter regions of several genes related to PS, such as *bch, crt* and *puc*, generally blocking their transcription [14], [15].

Data suggests there is a great divergence on the effects and the mechanisms of PpsR on PS regulation among different species of PNSB. While some PpsR proteins can directly sense redox signals and elicit a transcriptional response, such as in *Rubrivivax gelatinosus* [16] and *Rhodospseudomonas palustris* [17], others are constitutively prone to inhibit (or even promote) gene transcription until they interact with a second protein capable of binding to PpsR upon detecting an environmental/intracellular cue, as is the case of *Rhodobacter capsulatus* and *Cereibacter sphaeroides* [18]. These sensor proteins are AerR (<u>aer</u>obic repressor) [19], or its homologus PpaA (<u>p</u>hotopigment and *<u>p</u>uc* <u>a</u>ctivation) [14], [20], and AppA (<u>a</u>ctivation of <u>p</u>hotopigment and <u>p</u>uc expression, specific from *C. sphaeroides*) [21]. They can interact with PpsR, with a redox signal or light being their inducers. PpaA/AerR share the presence of at least one cobalamin binding domain. Instead of this domain, AppA contains a heme-binding domain designated SCHIC (<u>s</u>ensor <u>c</u>ontaining <u>ha</u>em instead of <u>c</u>obalamin) which belongs to the vitamin B12-binding domain family [22], and it works as an integration point for both redox and light inputs [18], [23], [24]. The best characterized PpaA/AerR anti-repressors have been demonstrated to bind with high affinity to cobalamin and even though their amino acid sequence is poorly conserved, their role as PpsR partner, as well as the proximity of the encoding genes in the chromosome, is a common feature in many PNSB.

*Rhodospirillum rubrum* is a PNSB which possess the aforementioned versatility of this group of bacteria. Moreover, it has been crucial in the elucidation of the basic fundaments of polyhydroxyalkanoates (PHA) metabolism [25]–[27] showing recently a great potential for the production of poly(3-hydroxybutyrate-co-3-hydroxyvalerate) (PHBV) from fructose in anaerobic conditions [28]. It has also been proposed as a biocatalyst for the production of hydrogen [29], [30], [31], pigments and vitamins [32]. In this microorganism, PS regulation shows particular features compared to other PNSB. For example, even though PpsR (codified in Rru_A0626) can be found in one of its three PS gene clusters (PGC) [33]–[35], nor RegBA/PrrBA neither PpaA/AerR or AppA homologous have been described in this species yet. Understanding how PS is regulated in *R. rubrum* might provide powerful tools for biotechnological exploitation of this microorganism as a source of industrially relevant bioproducts.

In this work, we characterized a spontaneous mutant of *R. rubrum* S1 (Rr02_01) with an impaired capacity of producing pigments in microaerobiosis. We made massive sequencing of Rr02_01 and found a disabling mutation in locus Rru_A0625, located in one of its three PGC. This mutation proved to have effects not only on PS related genes, but in many other biological processes, such as nitrogen metabolism, regulatory genes, and stress response. We propose HP1 as a fundamental protein for PS regulation in *R. rubrum*. Even though it has low identity with PpaA/AerR proteins, due to its role in PS activation and its proximity with PpsR, the here so-called HP1 can be considered the PpaA homologue in *R. rubrum*. However, it shows some remarkable features: It is the smallest version of the PpsR/AerR anti-repressor family and, more interestingly, possess no recognizable cobalamin binding domain.

## MATERIALS AND METHODS

### Growth conditions and plasmids

The strain used in this study was *R. rubrum* S1 (ATCC 11170, DSMZ 467T). It was stored in Luria-Bertani (LB) broth [36] containing 15% v/v glycerol at -80°C. To recover the strain, LB fructose (10 mM) was used. For the experiments described in the Results section, a modified RRNCO medium [37] containing K_2_HPO_4_ (19.1 mM) and yeast extract (1 g·l^-1^) (pH of 7.0) was employed. Precultures were grown in 25 ml of LB in closed 50 ml falcon tubes and incubated in the dark for 24 hours at 30°C and 200 rpm. Experimental cultures were performed in serum bottles sealed with cotton plugs or 20 mm-thick chlorobutyl plugs (Wheaton® W224100-202), with a starting OD_660_ of 0.05. The cultures were incubated at 30°C and agitated at 100 or 200 rpm, depending on the experimental conditions. A diagram of the entire experimental setup is illustrated in Fig. S1 [28]. For anaerobic cultures, bottles half-filled with medium were heated (70-80°C) before autoclaving using a microwave oven and degassed by injecting N_2_ for 15-20 min. After sterilization, phosphate (19.1 mM), fructose (13.3 mM), and the reducing agent sodium sulphate (0.01%) were added with a syringe to the bottles. Photoheterotrophic cultures were grown at 30ºC with 200 rpm of orbital agitation and an illumination intensity of 1.5 Klux.

Plasmids used in this work are listed in Table S1.

### Spectrophotometric techniques

Cell growth was monitored through the turbidity (OD_660_) of the cultures with a 96-well plate MultiskanSky spectrophotometer (Thermo Fisher Scientific®). Values of OD_660_ shown in this work were corrected considering the optical path length. The levels of the complex antenna, referred to as photo-membrane production (PMP), were estimated by normalizing the optical density at 880 nm by the OD_660_ (OD_880nm_ / OD_660nm_) using the same equipment. The ΔPMP is calculated as follows: ΔPMP = PMP_i_-PMP_c_, where PMP_i_ corresponds to the unknown sample and PMP_c_ refers to the corresponding negative control (unpigmented cells).

### Construction of *R. rubrum* deletion mutants

Gene deletion was achieved through homologous recombination using plasmid pK18msg, a variation of the mobilizable plasmid pK18*mobsac*B [38]. In pK18msg, the original multiple cloning site (MCS) was replaced by a new one containing the recognition site of type IIs restriction enzymes BsaI, BbsI and AarI. The replacement was done by means of EcoRI and HindIII restriction sites placed on the extremes (Table S2). This new plasmid allowed the efficient cloning of homologues regions via Golden Gate reaction as described previously [39], which combines ligation and the selected type IIs restriction enzyme in a single reaction pot. Homologues regions for Rru_A0625 deletion were amplified by PCR using the appropriate oligos (Table S2), obtaining the plasmid pK18_ΔA0625. *E. coli* MFD strain (auxotrophic for diaminopimelic acid, or DAP) was transformed by electroporation with each plasmid and used to conjugate *R. rubrum* wild type strain.

### Gene cloning for complementation experiments

For the creation of the merodiploid containing the wild type Rru_A0625 allele, the genomic region from the wild type strain was amplified by PCR with primers O_77 and O_78. The amplicons (∼1.8 kb), containing the restriction site for BamHI and SalI, were cloned in pSEVA231, giving pSEVA_HP1. For the complementation of strain ΔA0625 with the synthetic genes containing loci Rru_A0625 or Rru_A0625b, the plasmid pVSOP was used, amplifying the corresponding coding sequences with primers shown in Table S1. Plasmid pVSOP was constructed using parts and plasmids from Golden Standard assembly kit [40], although some plasmids form Marillonet collection [41] were also used. First, an inducible transcription unit (TU) was created. This TU contained promoter Pm (the 3-metyl-beanzoate (3MB) inducible promoter), BCD2 bicistronic ribosome binding site (RBS), a cloning site defined by SapI recognition sequence surrounding a differentiation cassette (DC), and *rpoC* transcriptional terminator. All the parts belonged to the Golden Standard kit with the exception of the DC. DC was composed of genes *crtE, crtY, crtI and crtB* from *Pantoea ananatis* necessary for biosynthesis of β-carotene. These genes were obtained by PCR using primers O_167 and O_168 and pAGM4673 as template. SapI restriction sites were designed so as to expose ATG and GGT codons after digestion, being both the cohesive extremes for the in-frame cloning of the coding sequence. This part, equivalent to a coding sequence part, was cloned into plasmid pICH41308 using BpiI. The promoter, RBS, DC, and terminator were assembled into plasmid pICH47742 using BsaI, giving pL1_VSOP. This construct together with XylS gene, were cloned into plasmid v07 by means of BpiI, resulting in plasmid pVSOP. All BpiI and BsaI digestions were performed as Golden Gate reactions, containing ligase T4 (NEB), and its reaction buffer, in the same pot, using the following incubation program: [(37ºC, 10 min; 16ºC, 10 min) x 4 times]; 37ºC, 10 min; 65ºC, 10 min.

### DNA transfer methods

For the preparation of chemocompetent, *E. coli* DH5α cells were grown in LB medium at 37ºC with agitation at 200 rpm until reaching an OD_660_ of 0.6. The protocol described elsewhere was followed [42].

*R. rubrum* cells were transformed using the electroporation method. To prepare the cells, a culture was grown overnight in LB-fructose medium (25 mM) in a half-filled 50 ml Falcon tube. The cells were then washed three times with 25 ml of a sucrose solution (0.3 M) that had been previously chilled on ice. After the third wash, the cells were resuspended in 200-300 µL of the same sucrose solution and 50 µL of the suspension was put into 0.1 cm electroporation cuvettes (1652083, Bio-Rad, Inc). Next, 0.5-2.0 µL of plasmid DNA was added to the cells. For the electroporation step, the cuvette containing the cell/DNA mixture was inserted into the Bio-Rad GenePulser electroporator (Bio-Rad, Inc), and subjected to an electrical sock using the following conditions: 1.8 kV, 200 Ω, and 25 μF. Immediately after the pulse, 500 µL of chilled SOC medium were added to the cuvette, and the mixture was allowed to rest for 10 minutes at room temperature. Then, the contents of the cuvette were gently transferred to a 1.5 ml Eppendorf tube to recover the cells, which were incubated at 30ºC and 200 rpm for at least 6 hours. After the recovery period, the cells were pelleted and plated onto LB agar plates containing the appropriate antibiotic for selection.

For conjugation, *E. coli* MFD/pK18_ΔA0625, and *R. rubrum* wild type were grown over night (ON) in LB (DAP 30 µM) and LB (fructose 10 mM) respectively. Cells were harvested and washed twice with saline solution (SS, NaCl 8.5 g·L^-1^) after which they were resuspended to reach an OD_600_ of 1. Subsequently, 500 µL of *R. rubrum* resuspension were mixed with 500 µL of MFD/pK18_ΔA0625. The mixture was centrifugated and the pellet was resuspended with 50 µL of SS. The cell mixtures were placed over a 0.22 µm pore MCE-filter (GSWP02500, MF-Millipore) on LB-agar plates containing 30 µM DAP and incubated statically ON at 30ºC. The biomass over the filters was resuspended in SS (1 ml), concentrated to100 µL by centrifugation (2600 rcf, 5 min) and plated on LB-agar with kanamycin (50 µg·ml^-1^). This procedure allows the selection of *R. rubrum* transconjugants which had performed the first homologous recombination. The resulting recombinant strains were verified via PCR. Five positives clones of each construct were grown ON in LB-fructose (10 mM) without antibiotic. Double-recombinant clones were selected by plating 100 µL of these cultures on RRNCO agar plates containing fructose (10 mM) and sucrose (5%). Several isolated colonies were transferred in parallel to LB and LB-km agar plates. The second recombination event was verified by PCR on sucrose resistant and Km sensitive clones.

### Analysis *in silico* of HP1

The protein structure prediction was performed with AlphaFold2 machine-learning algorithm [43]. The amino acid sequences were aligned employing the ClustalW algorithm in the MEGA11 software [44], applying the Gonnet protein weight matrix with a gap creation and gap extension penalty of 10 and 0.2, respectively. The conserved motifs of the proteins were explored by means of NCBI’s Conserved Motives browser. The percentage of the secondary structures of the proteins were determined using the Secondary Structure Server (https://2struccompare.cryst.bbk.ac.uk/). The Uniprot accession number of the proteins used for the analysis are mentioned in Fig S2.

### Transcriptomic analysis

For extracting the RNA, the wild type and ΔA0625 strains were grown on modified RRNCO medium with fructose (13.3 mM) under C2 (microaerobiosis) experimental conditions. The cultures were harvested during mid-exponential phase after the wild type strain had already initiated pigment production (aprox. 72h of incubation). The resulting pellets were immediately frozen and stored at -80ºC. RNA extraction was performed employing the High Pure RNA Isolation Kit (Roche), with an additional DNA digestion step that involved the addition of DNAse (Ambion™) and incubation of the mixture at 37ºC for 60 min. The purity, concentration, and integrity of the RNA samples were assessed using a Bioanalyzer2100 (Agilent Technologies, Inc). The integrity of the samples had an RNA Integrity Number (RIN) above 8.2 and a minimum concentration of 200 ng·mL^-1^.

The transcriptome resequencing of the RNA samples and the bioinformatic analysis were performed by Macrogen (Korea). Briefly, the ribosomal RNA was removed from the samples with the NEB Next rRNA Depletion kit, after that, the RNA was randomly fragmented and purified for short read sequencing and transformed into cDNA through reverse transcription process. The library construction was made using the TruSeq Stranded Total RNA Library Prep Gold Kit. The amplification of the fragments was made by PCR and the template size distribution was checked by Agilent Technologies 2100 Bioanalyzer using a DNA 1000 chip. The fragments with inserts between 200-400 pb were selected for the paired-end sequencing performed by NovaSeq 6000 Sequencing System. After sequencing, the quality of produced data was determined by the phred quality score at each cycle employing the software FastQC and trimmed with Trimmomatic. Trimmed reads were aligned against the genome sequence of *R. rubrum* S1 (accession number ASM1913455v1) through Bowtie2. Statistical analysis was performed calculating the Fold Change and doing nbinomWaldTest using DESeq2 per comparison pair. The significant results were selected on conditions of |fc|>=2 & nbinomWaldTest raw p-value<0.05. To ensure a more stringent comparison, only genes with a Benjamini-Hochberg (BH) p-value < 0.05 were subjected to further analysis. However, as an exception, 18 genes with a higher BH p-value were still included in the analysis if they were part of a transcription cluster (TC), that contained one or more genes with a significant BH p-value. In this work, a TC is defined as the group of genes codified in the same strand, separated by less than 250 pb, and either all up-regulated or down-regulated in the mutant strain. The definition of TC differs from that of operon in that we cannot assure that genes belonging to the same TC are expressed in a polycistronic RNA, as in the case of operons, but they are closely related in their transcription dynamic in the experimental conditions of the present study. RNA-seq data was deposited in the NCBI Reference Sequence Database under the accession number PRJNA940742.

## RESULTS

### Detection and characterization of the spontaneous mutant strain Rr02_01

Laboratory strains can adapt to anthropogenic conditions to which they are exposed during routine handling and successive passages. This uncontrolled evolutionary process may lead to silent mutations that are not easily perceived, or by the contrary, it can hit on genes responsible for clear and naked-eye detectable phenotypes. This is the case of the strain Rr02_01 isolated in our lab, derived from strain S1 of *R. rubrum*. Though expressing wild type behaviour during aerobic growth in rich media (LB), it manifested an impaired pigment production when cultured in solid medium. To assess whether this phenotype responded to oxygen tensions, we cultured both the wildtype and the spontaneous mutant (Rr02_01) strains in different regimes of aeration as detailed in the Materials and Methods section (see also Fig. S1).

In terms of biomass formation (OD_660_) no important differences were observed in the opened systems (O1, O2 and O3) where the gas in the headspace could be exchanged with the environment (Fig 1A). Nor was the case of C1, the most aerated of the closed systems. However, subtle but significant differences between both strains were observed in levels C2 and C3, considered to be least aerobic of all. In these two cases, wildtype formed more biomass than Rr02_01. As expected, the differential behaviour of both strains was regarding pigmentation. The OD_880_/OD_660_ ratio is directly related to the levels of photomembrane production (PMP), as it normalizes the amount of pigments (estimated throw the OD_880_) per unit of biomass (OD_660_). Considering the ΔPMP (the difference in the PMP of certain condition minus the basal level obtained in aerobic growth, i.e. O1), it can be easily observed that Rr02_01 does not produce pigments in any of the studied conditions (Fig 1B). These results suggest that the function failing in Rr02_01 is quite specific for sensing the redox state (directly or indirectly) and it is implicated in pigment synthesis.

**Fig. 1.**
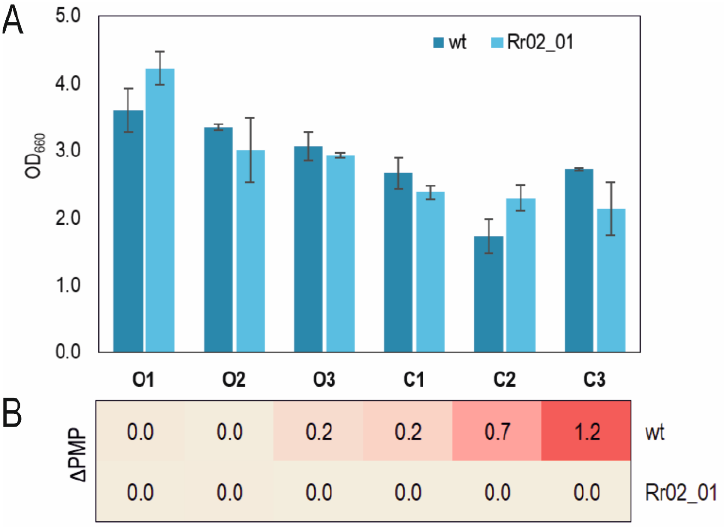
Effects of aeration level on biomass and pigment formation in the wild type and mutant (Rr02_01) strains. Different aeration levels were obtained controlling the agitation, the filling volume of the flask and the sealing of the bottles. (A) No relevant differences were observed in the final OD regardless of the aeration condition. (B) The wild type strain was able to pigment in every condition except of O1 and O2, whereas the spontaneous mutant strain did not pigment in any of them.

### The principal responsible of Rr02_01 inability to produce PM in microaerobiosis is a mutation in Rru_A0625 locus

To determine the genetic differences between the wildtype and RR02_01 strains, both genomes were sequenced. The reads were aligned using the reference genome of *R. rubrum* S1: NC_007643 and NC_007641 (chromosome and plasmid, resp.). After filtering mutations with low coverage (< 10) and high variation frequency (> 60%), a total of 2 unique differences present in RR02_01, but not in WT, were found. One of these differences was in locus Rru_A0625, placed in one of the three photosynthesis gene cluster (PGC). The second genetic variation was found in a gene that encodes a protein with a J domain (Rru_A0205), apparently not related to Rr02_01 unpigmented phenotype, which made us exclude locus Rru_A0205 from further analysis. The mutation in Rru_A0625 consists in a punctual deletion of a guanosine within a sequence of 7 consecutive G, leading to a frame shift that generates a STOP codon 13 nt upstream, and concomitantly giving a protein 60% shorter than the original. We will hereafter refer to this mutation as 7G6.

Locus Rru_A0625 is annotated as a 546 nt Open Reading Frame (ORF) predicted to codify a hypothetical protein of 181 amino acids and 20.25 kDa (here on called HP1). However, an in-frame ATG 111 nt upstream the predicted by the automatic annotation, was initially considered as the start codon. This was decided to avoid a possible underestimation of the correct size of the gene due to the lack of similar characterized genes in the database that could exclude an important portion of the resulting protein.

To prove that 7G6 mutation was related to pigment synthesis disability in Rr02_01, we knocked Rru_A0625 out in the wild type strain using the double recombination method (see Material & Method section). The resulting strain Rr02_05 (hereafter called ΔA0625) was unable to produce pigments in microaerobic cultures, such as the case of Rr02_01. When this strain was transformed with plasmid pSEVA_HP1 (containing wild type locus Rru_A0625 plus its 0.40 kb upstream and 0.75 kb downstream regions) the pigmented phenotype was recovered (Fig. 2). The strain ΔA0625 transformed with the empty vector pSEVA231 did not pigment.

**Fig. 2:**
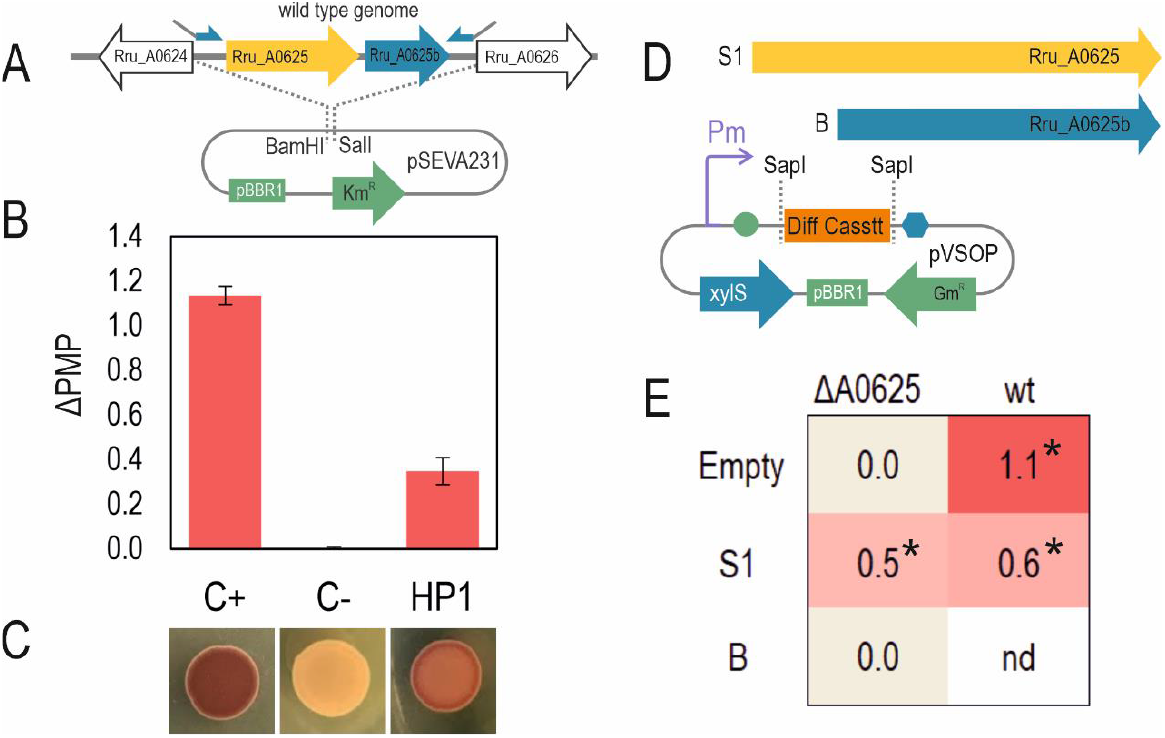
Complementation of strain ΔA0625. (A) The chromosomic region where the mutation 7G6 was found, was amplified using DNA from wildtype strain as template, and cloned into pSEVA231, giving place to pSEVA_HP1. The negative control was the empty vector pSEVA231. (B) Only pSEVA_HP1 could partially complement Rru_A0625 deletion in microaerobiosis (condition C2) using liquid modified RRNCO medium with fructose as carbon source. (C) In parallel, a drop (10 µl) of each preculture was incubated on solid LB-agar with fructose with equivalent results. (D) Rru_A0625 and Rru_A0625b were amplified by PCR and cloned separately into pVSOP, where the expression is controlled by the Pm-XylS system. (E) pVSOP_A0625^S1^, but not pVSOP_A0625^B^, was capable of complementing the deletion of Rru_A0625. The former plasmid, in the wildtype strain, caused a detriment in pigment formation respect to the empty vector in the same genetic background. Asterisks show significance regarding the negative control ΔA0625/pVSOP_Empty.

To confirm that Rru_A0625 locus and no other region in the chromosomic fragment cloned in pSEVA_HP1 complemented ΔA0625 phenotype, Rru_A0625 was amplified and cloned under the expression of Pm-xylS induction system in pVSOP vector (pVSOP_A0625^S1^) (Fig 2D). Another ORF (here called Rru_A0625b), besides Rru_A0625, can be predicted in the vast intergenic region between the latter and the next locus, Rru_A0626. A plasmid containing this ORF was also constructed (pVSOP_A0625^B^). The strain carrying pVSOP_A0625^S1^, was the only one capable of pigmenting under microaerobic conditions, meanwhile pVSOP_A0625^B^, was not (Fig. 2E). This meant that undoubtedly, Rru_A0625 malfunctioning was the responsible for Rr02_01 and ΔA0625 non-pigmented phenotypes.

### *In silico* analysis of Rru_A0625 gene product

HP1 exhibits a certain degree of similarity with proteins containing the Cobalamin-B12 binding (CB12B) motif (Fig 3A). Among these proteins, the one with the highest identity to HP1 is a hypothetical protein from *Novosphingobium* sp. Gsoil 351, with a sequence identity of 45% (62% coverage). The conserved motif of CB12B is a common feature among other well-known PpsR antirepressors belonging to the PpaA/AerR family, generally placed in a similar position relative to *ppsR* in different species. When HP1 is aligned with some of these proteins, it can be seen that HP1 lacks most of the conserved amino acids including those of the CB12B motif (Fig. S2). Besides, with only 181 amino acids, HP1 is 17% smaller than PpaA from *Rb. Capsulatus*, the smallest experimentally tested member of this family.

**Fig 3.**
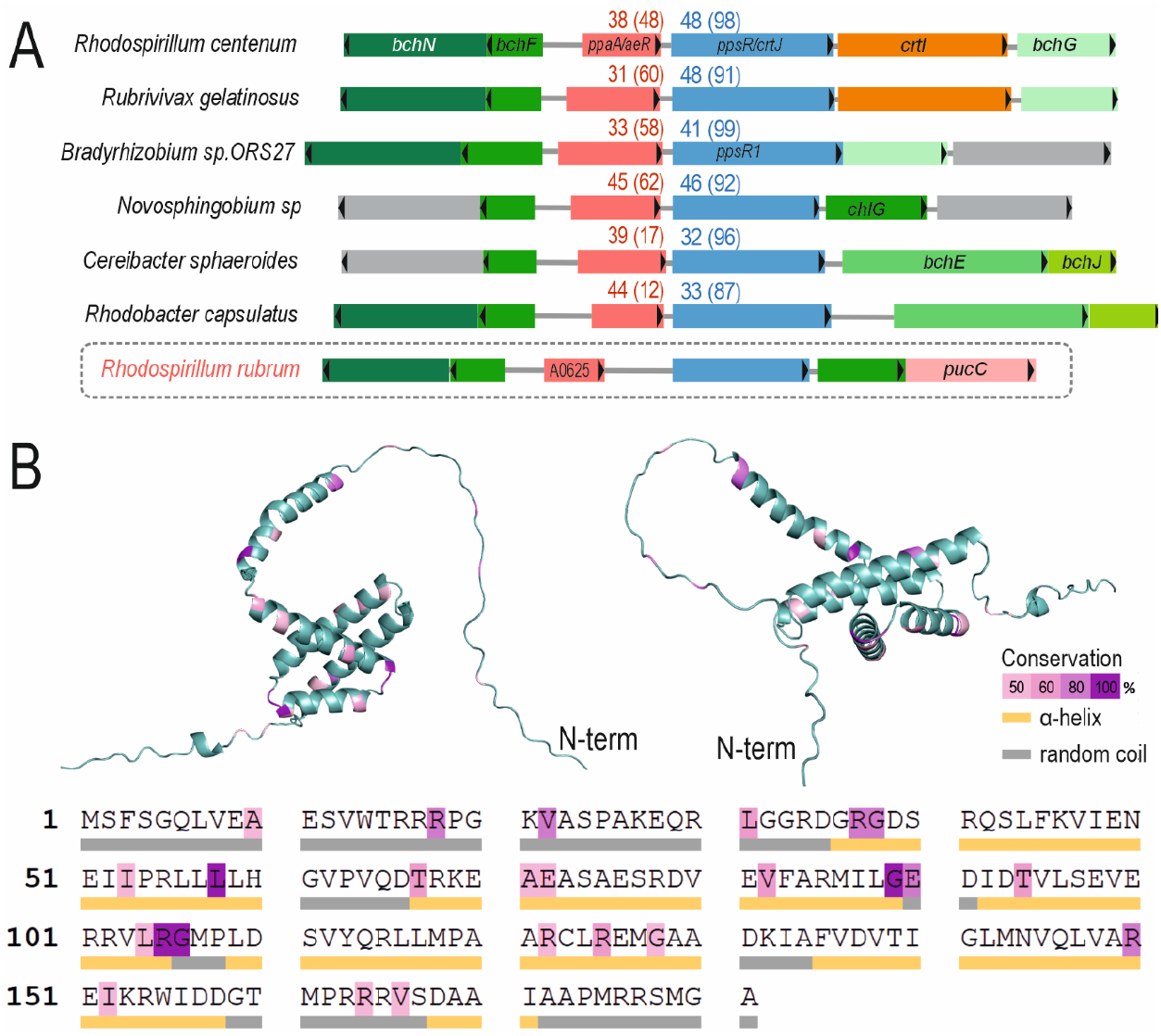
In silico analysis of HP1. (A) Scheme of part of the photosynthetic gene cluster that contains *ppsR*. The transcription sense is indicated with a black triangle in each coding sequence. The percentages of identity and coverage (in brackets) of *ppaA/aerR* and *ppsR* with the corresponding genes in *R. rubrum*, are presented in red and blue respectively. Homologous genes are depicted with the same colour. Genes with unknown function are coloured in grey. The analysis was performed with BLASTp. (B) 3D-structure of HP1 was predicted using AlphaFold2. The colour code indicates the conservation degree for each amino acid (pink to dark violet), and the secondary structure.

The secondary structure also shows HP1 is highly disordered, with only 50% of its sequence predicted to have α-helix structure. It contains only one cysteine in the position 124. These features together, make HP1 from *R. rubrum* stand out in relation to the other members of the PpaA/AerR family.

### Regulatory redundancy: PSA is controlled by more than one regulator and more than one cue

Previously, Rru_A0625 function was tested in aerobic-microaerobic conditions in the absence of light. In other species, such as *C. sphaeroides*, pigmentation is also activated by light [18], [24], [45]. Then, we cultured the wildtype and knock out (KO) strains in anaerobic conditions and RRNCO-fructose medium (condition AN, Fig S1), under the exposure of light to see the effects on growth and pigment synthesis. Bicarbonate (NaHCO_3_) was added as a source of CO_2_, a fundamental electron sink to promote growth in anaerobiosis. We also tested the effect of an alternative electron acceptor, the DMSO (Fig 4). This compound can capture electrons coming from the respiratory chain or the photosynthesis by means of a DMSO-reductase [46], indicating the possible implication of these electron chains in the process of study.

**Fig. 4:**
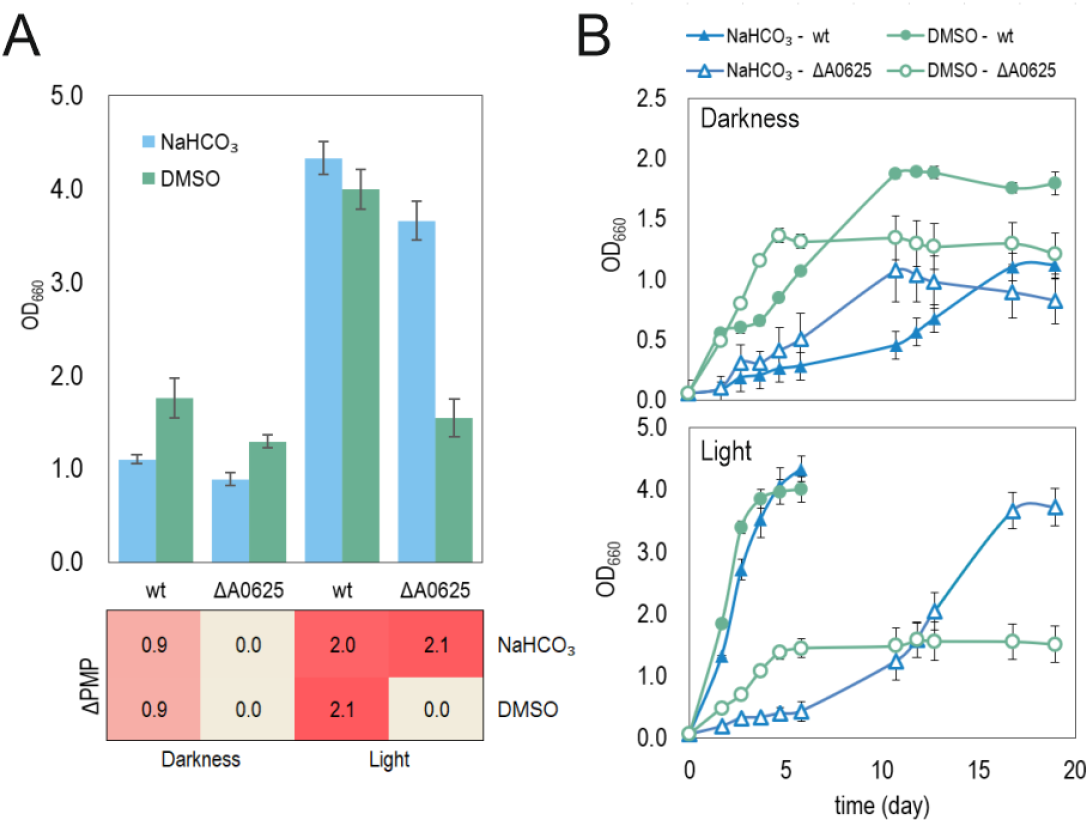
Growth and PSA activation in anaerobic conditions. Strains ΔA0625 and wild type (wt) were grown in RRNCO medium with fructose under anaerobic conditions in the presence or not of light. Bicarbonate (NaHCO_3_) was added to favour growth in the absence of oxygen as an electron sink. DMSO (an alternative electron acceptor) was also used to infer if the electron chain was implicated in the signalling process. Cultures were grown anaerobically (AN) with light or in darkness. The (A) final OD_660_ / ΔPMP and the (B) growth curves are shown in this figure.

In photoheterotrophic conditions, both strains were able to grow and develop the PSA. However, strain A0625 had a longer lag-phase. In darkness, the anaerobic culture of the KO mutant strain did not pigment, contrary to wildtype strain. Both the wildtype and mutant strains grew better with light. The addition of DMSO annulled the light-induced pigmentation only in the KO strain. These results together prove there is an additional system in *R. rubrum* other than Rru_A0625 that promotes the synthesis of PSA responding to light, probably by means of a cytochrome placed under the quinone pool, from where the DMSO-reductase takes the electrons to reduce DMSO to DMS [46].

### Transcriptomic of ΔA0625 mutant revels pleiotropic effects on cell physiology of HP1

A clear impact on pigment production can be attributed to Rru_A0625 malfunctioning. To address if its effects extend beyond the expression of the photosynthesis apparatus, we performed a RNAseq experiment on cultures from both strains (wildtype and Δ0625) grown in RRNCO mineral medium with fructose as carbon source and microaerobic conditions (C2). The comparison was made in a timepoint carefully selected to study the cells in a clearly differentiated stage, i.e. in the exponential phase after the wildtype strain had started to produce pigments (day 3 of culture).

A total of 273 genes were affected after Rru_A0625 deletion, 7.4 % of the total number of genes in *R. rubrum*. From these genes, 227 (83%) were downregulated in strain Δ0625, while 47 (17%) of them were upregulated. Almost half of the affected genes (153 genes, 56%) were grouped in 52 transcription clusters (TC definition is in the Material & Method section). A Clusters of Orthologous Groups (COGs) analysis of the affected genes (Fig. S3) revelled the multiplicity of processes affected by deletion of Rru_A0625 locus. To have a clearer picture of biological role of HP1, the list of affected ORFs where manually re-grouped according to the biological process they could be involved in (Photosynthetic Apparatus or PSA, Regulation, Sugar and Polysaccaride Metabolism, Stress Response, Amino acid Metabolism, Nitrogen Metabolism, etc) (Fig. 5). When the biological process of certain gene was not clear, it was classified as unknown.

**Fig. 5:**
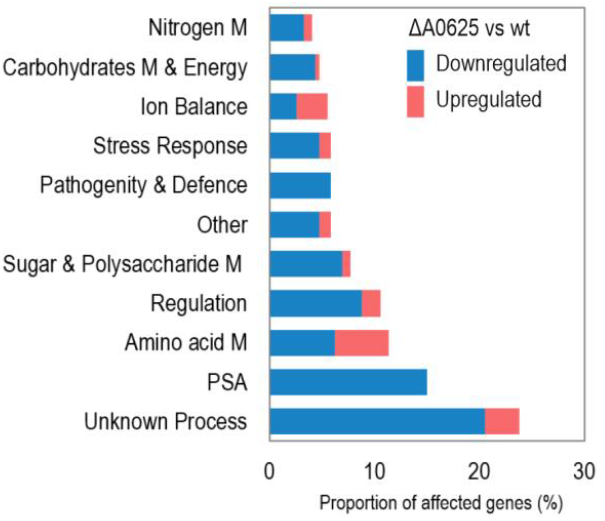
Transcriptomic impact of Rru_A0625 mutation. The initial COG classification was further refined through manual re-elaboration and simplification, whereby genes were grouped based on their biological processes. Data from the literature and/or the *in silico* analysis of their aminoacidic products was used to cure the classification. The proportion of genes is expressed relative to the total affected genes. M, metabolism; PSA, photosynthetic apparatus related genes.

**PSA**. most genes grouped in the structural category, participate in the PSA formation. In fact, this category accounted for a 12% of all the affected genes in terms of the physiological process they are involved in, preceded only by genes with unknown function. All the genes classified as PSA were negatively affected by Rru_A0625 deletion, indicating the clear positive effect that HP1 has over this process. The two largest transcription clusters (TC) found in this transcriptomic analysis, contained 12 and 10 genes almost exclusively related to PSA, with an average fc of -23 and -83 respectively, by far the transcription rates most affected by Rru_A0265 deletion, as expected considering the observable phenotypic changes in both strains.

PNSB generally encodes PSA components on a discreet region of the chromosome, the PGC, which includes genes related to the synthesis of bacteriochlorophyll, carotenoid, reaction center (*puf*), light-harvesting complexes (*puh*) and regulatory proteins (PpsR) [35]. In *R. rubrum* the PGC is divided in three: PGC 1, 2 and 3. HP1 and PpsR are located in the second PGC (PGC 2) where the highest fc (−44) corresponds to *bchH* (Rru_A0621), a magnesium chelatase. However, both HP1 and PpsR are codified in the opposite strand, at the 3’ end of a second transcription cluster (TC) with a lower average fc than the one containing *bchH* (−23 vs -4) (Fig R7). Although it is difficult to make an accurate estimation on the transcription level of Rru_A0625, since it was deleted from strain ΔA0625, we can confirm that *ppsR* is not significantly up- or down-regulated, indicating that its transcription is disconnected from its TC partners, and moreover, from the average transcription level of the whole PGC1 (−23). PGC 1 was the most fragmented in transcriptional terms, containing only one TC and three transcriptionally “isolated” genes. It is also the least down-regulated PGC in the mutant strain, with the highest fc corresponding to Rru_A0495 (fc = -7).

**Fig. R7:**
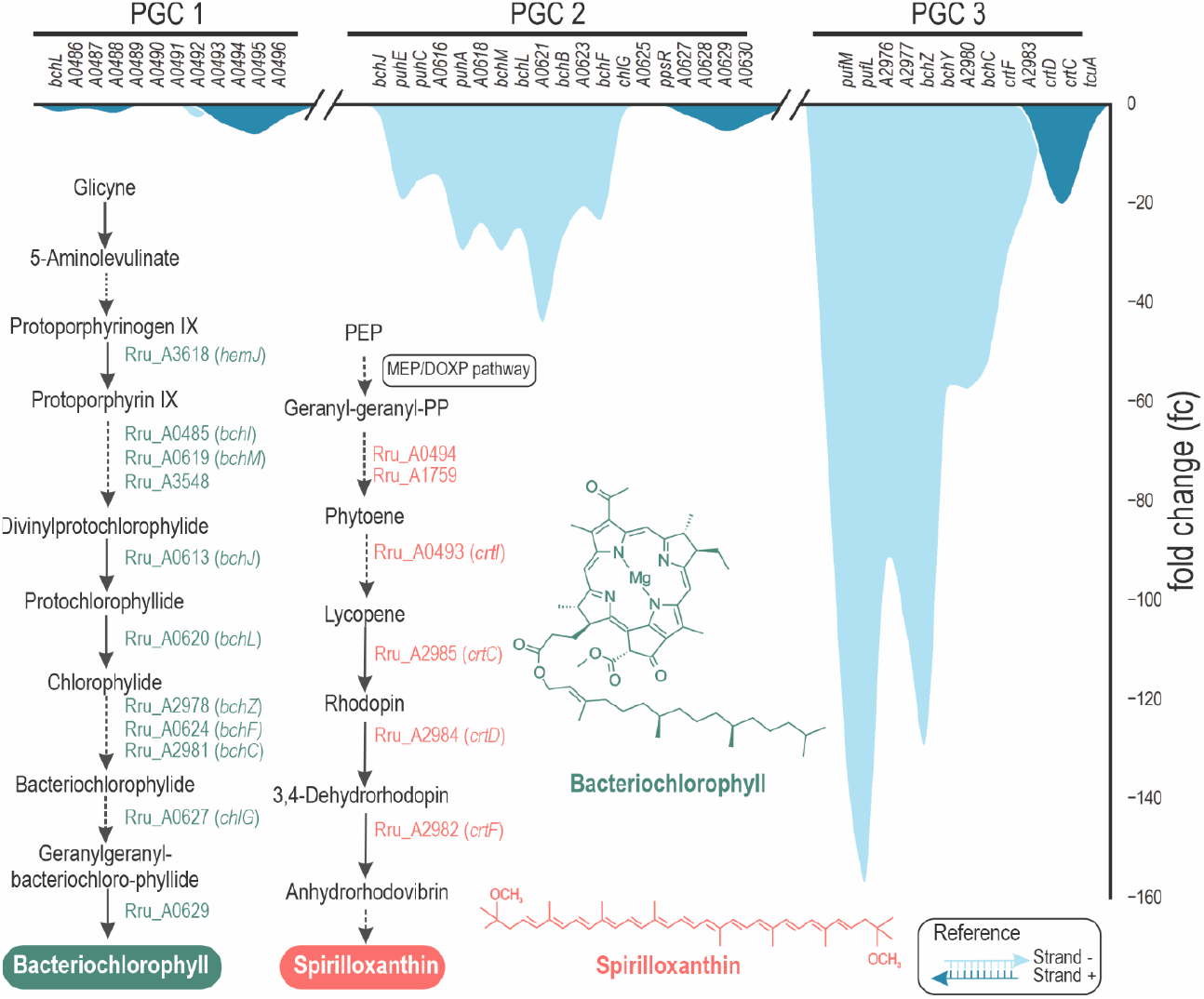
Fold change expression of the genes located in the three photosynthetic gene clusters (PGC). In *R. rubrum* PSA related genes are manly grouped in three clusters: PGC 1, 2 and 3. The most affected genes are placed in the PGC 3. The second most affected is the PGC 2, where HP1 is codified together with PpsR, in the positive strand. Interestingly, *ppsR* transcription is virtually unmodified in the mutant strain, placed in a fold change valley between two down-regulated TC. The PGC 1, on the other extreme, has the lowest fold change levels of the three PGC. Remarkably, in these regions, genes codified in the positive strand were less affected than those in the negative strand.

### Nitrogen Metabolism

After PSA, genes corresponding to nitrogen metabolism were the most affected in the mutant strain (Fig 8). Operon *nifHDK* codes for the nitrogenase complex, which is in charge of N_2_ fixation with the collateral production of H_2_ [47]. This complex is constituted by a dinitrogenase (MoFe protein or component I) and a nitrogenase reductase (Fe protein, or component II). These genes are 7 to 9-fold downregulated in strain ΔA0625. Elsewhere in the chromosome, a TC composed by *nifQ, fdxB, nifX, nifN* (Rru_A2281 to Rru_A2285), coding for accessory proteins associated to FeMo-co maturation, have a similar fc to *nifHDK* [48]. Surprisingly, other proteins required for this purpose in *Azotobacter vinelandii*, such as NifB (Rru_A0994), are not affected in ΔA0625 mutant.

**Fig. 8:**
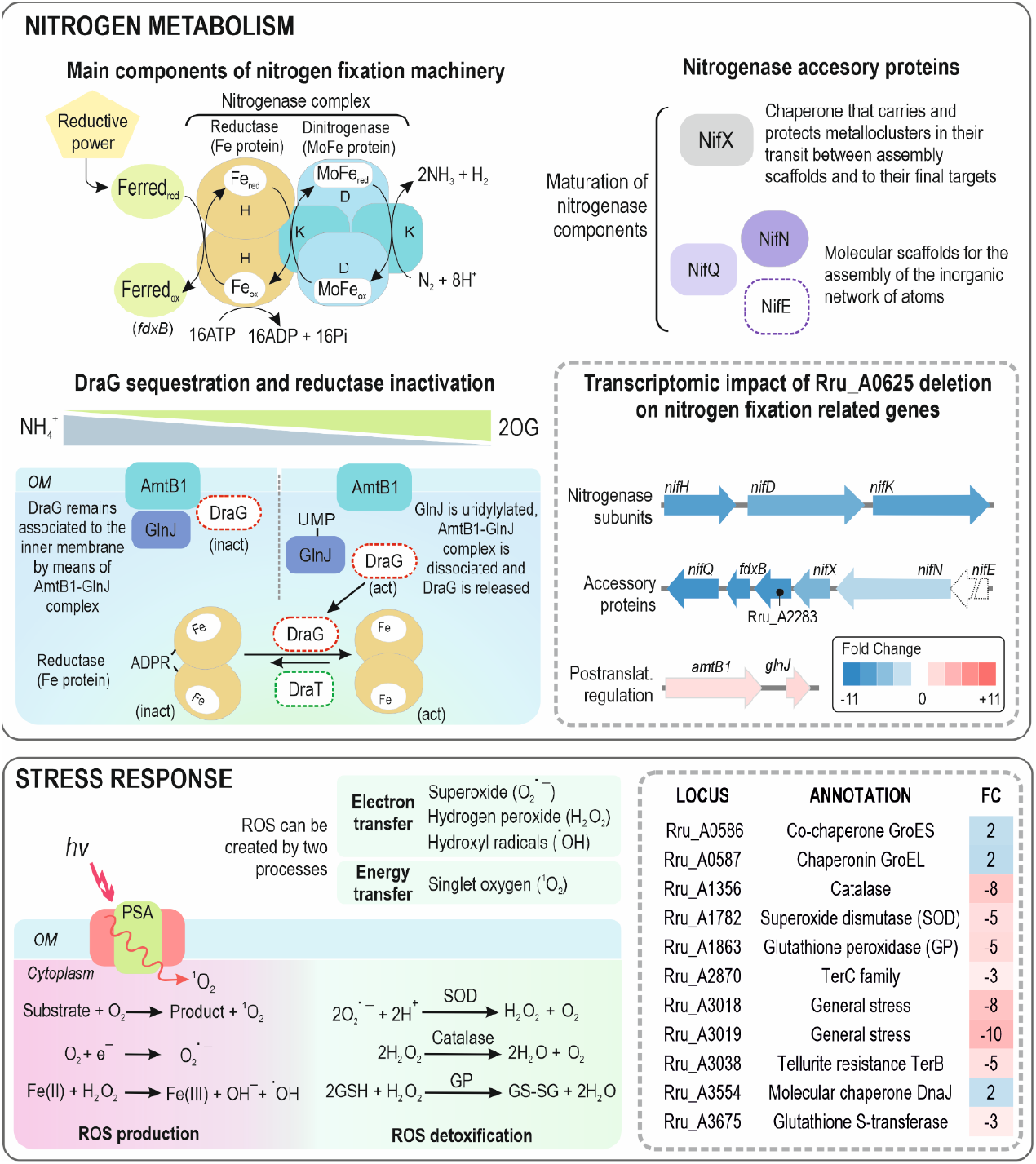
Impact of Rru_A0625 deletion on genes related to nitrogen metabolism and stress response. Nitrogenase subunits (*nifH, nifK* and *nifD*) and some accessory proteins were downregulated in ΔA0625 mutant. On the other side, *amtB* and *glnD*, implicated in the post-translational inhibition of the nitrogenase, were upregulated. Electrons to fix N_2_ are supplied by a ferredoxine (Ferred). *fdxB* product could serve as an electron donor to nitrogenase as it was the case of the cyanobacterium *Anabaena* sp. strain PCC 7120 [56]. The latter metallocluster-prosthetic group is assembled and completed by the sequential activities of several biosynthetic accessory proteins. Reductase activity is modulated by the action of two enzymes, DraG (dinitrogenase reductase activating glycohydrolase) and DraT (dinitrogenase reductase ADP-ribose transferase) that act in a reciprocal manner [57], [58]. AmtB1-GlnJ complex reversibly sequester DraG close to the outer membrane (OM), preventing it from acting [59]. Depending on the levels of ammonium and 2-oxoglutarate (2OG), GlnJ can be uridylylated, dismantling the complex and releasing DraG. Most of the genes coding stress responsive proteins, have a role in ROS detoxification. ROS is created in secondary reactions during metabolism by means of electron transfer or energy transfer. SOD, catalase, glutathione peroxidase (GP) dampen ROS side effects [60]. Other chaperone like proteins were also affected in ΔA0625 mutant. Elements which were not affected but appear in the scheme, are represented with a dotted line.

On the other side, two genes with significative fc are upregulated in ΔA0625 strain: *amtB1* (Rru_A1129) and GlnJ (Rru_A1130). The latter belongs to the P(II) protein family and represses nitrogen fixation by inhibiting different levels of its complex regulation [49], [50], [51]. Thus, it can be seen that HP1 mutation affects either directly or indirectly multiple targets of the nitrogen fixation machinery at the transcriptional level.

### Stress response

17 genes were included in this category. Some oxygen induced stress proteins seemed to be upregulated by HP1 (Table S3, Fig 8). For example, genes codifying for a catalase (Rru_A1356), a glutathione peroxidase (Rru_A1863), a superoxide dismutase (Rru_A1782) and a glutathione S-transferase (Rru_A3675) had a fc in the mutant strain of -8, -5, -5, and -3 resp. It was particularly striking the affection of two genes related to tellurite detoxification, one codifying a tellurite resistance TerB family protein (Rru_A3038) and other TerC (Rru_A2870), (fc of -5 and -3, respect). Only three genes showed higher transcription rates (2-fold) in the deletion strain compared to the wt: a molecular chaperone DnaJ (Rru_A3554), co-chaperone GroES (Rru_A0586) and its TC partner, a chaperonin GroEL (Rru_A0587).

### Other biological processes affected by Rru_A0625 deletion

Important biological processes other that the aforementioned, were affected in ΔA0625 mutant.

The group of Amino acid Metabolism contains as many upregulated as downregulated genes. Loci Rru_A3466 (*rocD*) and Rru_A3467, localized in the same TC, are the most upregulated genes in the mutant strain (fc = 5 to 7). *rocD* contains an ornithine--oxo-acid transaminase related to the metabolism of arginine and proline, while Rru_A3467 product is an arginine deiminase (ADI). ADI is highly conserved in bacteria and it results in the conversion of arginine to ornithine, carbon dioxide and ammonia, with the concomitant release of ATP [52]. It is employed by many bacteria to buffer against acidic environments, raising intracellular pH [53]. *hutI* (Rru_A1303) and *hutU* (Rru_A1304) were also downregulated. Codified in the same TC, their transcription products are a urocanase and an imidazolonepropionase, which catalyse the second (EC 4.2.1.49) and third (EC 3.5.2.7) steps out of four in the universal histidine degradation pathway [54].

Internal polysaccharide storage and exopolysaccharide production represented 8% of the affected genes. A protein with an α-amilase domain (Rru_A2294) was 8-fold down-regulated in the absence of Rru_A0625. Another gene product affected (fc = -3) with this type of domain was GlgB (Rru_A2576), a glycogen branching enzyme. Besides, two glycogen debranching enzymes GlgX (Rru_A0505 and Rru_A2577) were -5 and -2 fold affected.

More relevant for the number of genes altered than for the magnitude of their fc, is the category of Regulation (9% of the total genes, or 25 genes), where most of the genes are up-regulated (80%), including a ChrR family anti-sigma-E factor, a LysR family transcriptional regulator and a sigma-70 family RNA polymerase sigma factor, all with a fc of -4. Interestingly, in *C. sphaeroides*, ChrR sequesters σ^E^ (also known as RpoE) which is implicated in stress response to ROS.

The Pathogenicity and Defence category contained genes manly related to secretion systems. The most affected was a protein that belongs to AI-2E family transporter (Rru_A3520, fc -6). TqsA has been shown to mediate transport of the quorum-sensing signal autoinducer 2 (AI-2) [55]. In the case of Ion Balance (5% of the total genes) half of them augmented their expression after Rru_A0625 mutation. Carbohydrate Metabolism and Energy accounted also for 5% of the total affected genes, being the most affected a NAD-dependent succinate-semialdehyde dehydrogenase (Rru_A0462) with a fc of -5.

## DISCUSSION

The starting point of this study was the spontaneous mutant strain Rr02_01 with impaired pigmentation. We could track the genetic origin of this abnormal phenotype by genome sequencing. Only two differences respect to the wild type strain were found. One was located in locus Rru_A2050, whose product is a J domain containing protein. This kind of proteins are co-chaperones capable of interacting with Hsp70 to assist homeostasis maintenance in the cell [61]. The other genetic difference was a single nucleotide deletion in locus Rru_A0625, located in one of the three so-called photosynthetic clusters (PGC). The deletion of Rru_A0625 in a wild type genetic context led to slightly pink colonies, reproducing Rr02_01 phenotype. Rru_A0625 translated into an enigmatic protein, which enclosed an apparent contradiction: it was somehow similar to proteins containing cobalamin-binding domains though this likeness was distorted enough to make any related domain (or any other present in databases) unrecognizable on its sequence or structure. It is annotated as a hypothetical protein with unknown function, reason why we called it HP1 along this work.

The relative position in *R. rubrum* brought another clue about its biological function. Immediately downstream Rru_A0625, the coding sequence of PpsR was found (Fig 3B). This protein has regulatory functions in many PNSB, principally as a repressor to avoid that photosynthesis reactivity jeopardises cell physiology through ROS formation in the presence of oxygen [9]. And, as is the case of many other species, the redox sensing task is outsourced to a second protein that, after being modified by its ligand, alters PpsR ability to repress photosynthetic genes. A diverse collection of proteins named PpaA, AerR and AppA, with a low degree of identity, controls PpsR activity in different species. It was reasonable to infer that both proteins (HP1 and PpsR), played a similar partnership in *R. rubrum*. Unfortunately, we were not able to obtain the single knock out in Rru_A0626 locus, even though we had succeeded to get the first recombination of the suicide plasmid pK18msg. Independently on the condition used to select the second recombination (aerobic with LB, anaerobic with RRNCO-fructose medium in light or darkness, with or without DMSO, etc), the results were negative (data not shown).

The transcriptomic analysis of the wild type and deletion mutant ΔA0625 grown micro-aerobically in the dark with fructose as carbon source, manifested the affection not only of the genes involved in pigment synthesis, but also other related to stress response, nitrogen fixation, amino acids, sugars and polysaccharide metabolism, and genes involved in gene/RNA/protein regulation, reinforcing the importance of coordinating these biological processes to respond efficiently to changes in the environment.

### Two distinct systems for controlling PSA synthesis

In darkness, *R. rubrum* is capable of assembling its photosynthetic apparatus (PSA) when oxygen levels are low enough even in the absence of light. We used closed systems to express PSA in microaerobic conditions to distinguish the incapacity of producing pigments of strain Rr02_01. When the cultures were anaerobic and cells were exposed to light, the null mutant ΔA0625 recovered the capacity of producing pigments but with a longer lag phase and slower growth. This indicates that a second system induced by light is involved in this adaptation, but less efficiently than HP1. HP1 is likely to work as a fast mechanism for microaerobic/anaerobic adaptation in response to a redox cue. Since the activation of PSA in the dark was not hampered by DMSO, it is possible that the input for HP1 activation does not come from the electron transport chain (ETC). However, it cannot be rule out a mechanism similar to that of PrrBA in *Rb. sphaeroides*, where the electron flow from *cbb*_*3*_ oxidase, and not the direct intervention of molecular oxygen, inhibits PrrB [46]. This oxidase functions independently of photosynthesis and is inoperative concurrently with the activation of PSA [62].

In darkness, DMSO had a greater impact on the growth dynamics than on pigment production. However, under light conditions, DMSO completely inhibited PSA formation in strain ΔA0625, with the expected negative effect on growth. Thus, the alternative regulatory mechanism may be activated by a light-dependent component downstream of the quinone pool, which is the preceding step to electron deviation produced by DMSO reductase. Nonetheless, this alternative signalling pathway appears to be redundant and dispensable, as the wild-type strain is able to grow both in the presence and absence of DMSO. This redundancy in PSA activation mechanisms could work as an emergency wheel in case HP1 cannot exert its function: even not as good (fast) as the regular wheel, it effectively enables to overcome the glitch.

The small size of HP1 (181 aa in its annotated version) compared to other similar proteins (218 aa from *Rb. capsulatus*) and the lack of conserved domains, let us think we are in front of the minimized version of a PpaA/AerR-like sensor, where the evolution has erased every known feature of the protein without extinguishing its sensing capability. It is unlikely that HP1 can bind either cobalamin or haem. The mechanism for sensing the redox state of the environment seems quite mysterious. In *Rb. capsulatus* and *Rb. sphaeroides*, an intramolecular disulphide bond plays a crucial role in PpsR activation. However, in *R. palustris* CGA009, PpsR2 lacks the ability to form an intramolecular disulphide bond since it has only one Cys residue. It has been proposed that the reduction of this Cys residue breaks an intermolecular disulphide bond necessary to change its activation state [17]. In a similar fashion, it could be possible that the single cys present in HP1 intervenes in an intermolecular association of HP1 required for transmitting the signal to PpsR or any other potential transcription factor.

### The diverse regulatory targets of HP1

As expected, the consequences of Rru_A0625 absence affected more deeply genes implicated in bacteriochlorophyll (*hem, bch*) and carotenoid (*crt*) synthesis. It is noteworthy that *ppsR* transcription is stable independently on the presence of HP1, in spite of being located immediately next to the second highest regulated TC. This is also the case of CrtJ (PpsR homologue in *Rb. capsulatus*) in a *ΔaerR* strain, whose expression level were comparable to those of the wild-type parental strain, indicating that deletion of *aerR* did not affect the cellular level of CrtJ [63]. PpsR2 in *Bradyrhizobium* sp shows the same constitutive expression regardless the aeration level [35]. These facts, manifests how tight the control on these regulatory components must be to avoid an excessive repression of the PSA.

After PSA related genes, nitrogen fixation genes were the most affected by Rru_A0625 deletion. This work confirms once more the imperative necessity of tuned coordination of the principal biological processes, such as photosynthesis, nitrogen fixation, and central metabolism. Our previous work [28] showed a remarkable role of the concerted action of CCB cycle, H_2_ evolution and specially PHBV synthesis for balancing redox poise. With an excess of ammonium sulphate and nitrogen coming from yeast extract (both at 1 g·l^-1^), nitrogen fixation does not seem to be crucial for survival, and so HP1-dependent transcription of nitogenase components *nifHDK* and *nifNX-fdxB-nifQ* in the wild type strain, does not appear to be attending such need. HP1 mission, in this case, appears to be the activation in low oxygen tensions of the burdensome nitrogen fixation machinery, probably to dissipate reducing power. AmtB seems to have two roles in the cell. The first function, is to act as a channel for uncharged ammonia. The second one is to interact with PII [64]. There are three PII homologs in *R. rubrum*: GlnB, GlnJ, and GlnK. All three PII homologs can regulate the activity of GlnE (ATase), which reversibly adenylylates glutamine synthetase [65]. GlnB and GlnJ, play a crucial role in controlling the reversible ADP ribosylation of dinitrogenase reductase catalyzed by the DraT/DraG (dinitrogenase reductase ADP-ribosyl transferase/dinitrogenase reductase-activating glycohydrolase) regulatory system [66]. GlnJ together with AmtB, can sequester DraG in the cytoplasmatic membrane, where DraG cannot act. Notably, ADP/ATP ratio and 2-KG disrupt this interaction, suggesting this is the way that GlnJ measures the cellular energy status [64].

Amino acid metabolism is not re-shaped towards a clear direction when HP1 is absent. However, HP1 seems to repress enzymes that could be involved in amino acid degradation of arginine, proline and histidine. Genes related to the stress response, on the contrary, seem to be summoned by HP1 to fight against oxidative damage, as it can be deduced by the positive effect of HP1 over the expression of catalase, glutathione peroxidase, superoxide dismutase enzymes. It is also interesting the positive effect that HP1 has on the transcription of two regulators that could be associated to stress response: ChrR anti-sigma factor (Rru_A0721) and a sigma-70 from the ECF subfamily (Rru_A3287). These sigma factors are responsible for orchestrating cellular responses to a wide range of environmental stimuli, including stress response. Typically, an ECF sigma factor is co-transcribed with a gene that encodes a negative regulator (anti-sigma factor). These anti-sigma factors act as inhibitors, regulating the activity of their cognate ECF sigma factors [67], [68]. The co-regulation of both antagonistic proteins, coded in different regions of the genome, raises the question of whether they interact in the same biological processes or have unrelated targets.

Related to internal polysaccharide metabolism, enzymes implicated in glycogen metabolism (a glycogen branching and debranching enzymes) were affected, showing that HP1 activate the expression of these enzymes together to maintain a balance between glycogen biosynthesis and degradation processes, as it has been already described [69].

Finally, the amount of the regulatory genes affected by HP1 is remarkable. Due to the amount of functions altered in strain ΔA0625, it is reasonable to think that many of them may not be directly controlled by HP1, but by a transcriptional regulator whose expression is directly or indirectly under HP1 control. This fact places our protein in a higher hierarchical position than expected.

## Conclusions

Despite the low similarity with other PpaA/AerR members, its role in PSA formation and its relative position to *ppsR* are solid arguments to include HP1 in this family of proteins. HP1 proved to be a key component in the regulation of photosynthesis. Its effects extend over genes implicated in a variety of biological processes, such as nitrogen fixation, stress response, amino acid metabolism, transcriptional/post-translational regulation, among others. We cannot exclude the possibility that this variation in the transcriptomic profile is not directly promoted by HP1, but other regulatory proteins intervene. Our protein reacts to changes in the redox state of the cell, as the vanguard mechanism for microaerobiosis/anaerobiosis adaptation. We propose to call HP1 PpaA (<u>p</u>hotopigment and *<u>p</u>uc* <u>a</u>ctivation), since it describes better HP1 function rather than AerR (<u>aer</u>obic repressor). This protein stands as an interesting regulator to be manipulated in different biotechnological approaches where PSA is dispensable, such as the production of hydrogen or polyhydroxyalkanoates in the absence of light.

## ACKNOWLEDGEMENTS

Authors want to acknowledge the financial support from the CSIC Interdisciplinary Thematic Platform (PTI+) Sustainable Plastics towards a Circular Economy (PTI-Susplast+), the Community of Madrid (P2018/NMT4389) and the Spanish Ministry of Science and Innovation under the research grant BIOCIR (PID2020-112766RB-C21). Santiago R. de Miguel is a recipient of a predoctoral FPI grant PRE2021-098281. Authors would also like to thank Ana Valencia for her constant help and technical assistance.

## AUTHOR CONTRIBUTIONS

MG conceived the presented idea and designed the experiments, supervised the work, and wrote the original draft. SM helped with the design of the experiments and conducted the RT-qPCR experiment and polymer measurements. AP obtained funds for the project, supervised the work, reviewed, and edited the manuscript. All authors discussed and contributed to the final version of the manuscript. All authors read and approved the final version of the manuscript.

